# DECREASE IN DNA METHYLATION 1-mediated epigenetic regulation maintains gene expression balance required for heterosis in *Arabidopsis thaliana*

**DOI:** 10.1101/2025.08.21.671646

**Authors:** Kodai Matsuo, Renmin Wu, Hiroto Yonechi, Terumi Murakami, Satoshi Takahashi, Asuka Kamio, Mst. Arjina Akter, Yoshiki Kamiya, Keita Nishimura, Takakazu Matsuura, Kaoru Tonosaki, Motoki Shimizu, Yoko Ikeda, Hisato Kobayashi, Motoaki Seki, Elizabeth S. Dennis, Ryo Fujimoto

## Abstract

Heterosis, or hybrid vigor, is characterized by superior phenotypic performance in F₁ hybrids compared to their parental lines, and its expression is influenced by both genetic and epigenetic factors. In this study, we investigated the role of epigenetic regulation, particularly DNA methylation mediated by DECREASE IN DNA METHYLATION 1 (DDM1), in early seedling biomass heterosis using hybrids between *Arabidopsis thaliana* accessions Columbia-0 and C24. Loss of DDM1 function in F₁ hybrids resulted in a significant reduction of rosette diameter, confirming that DDM1 is essential for heterosis. Transcriptomic and epigenomic analyses revealed extensive genotype-specific changes in gene expression and DNA methylation patterns in *ddm1* mutants. Notably, *ddm1*-F₁ hybrids exhibited upregulation of genes categorized into ‘xyloglucan:xyloglucosyl transferase activity’ and downregulation of genes categorized into ‘circadian rhythm’, which may contribute to reduced growth vigor. Whole-genome bisulfite sequencing showed widespread hypomethylation in *ddm1* mutants, yet the overlap between differentially methylated and expressed genes was limited, suggesting indirect or context-dependent regulatory effects. Additionally, we examined the role of salicylic acid (SA) in heterosis using SA-deficient mutants and found that variations in endogenous SA levels did not correlate with biomass heterosis under normal conditions. Our findings suggest that heterosis in *A. thaliana* is dependent on the maintenance of parental epigenetic divergence, particularly in DNA methylation patterns mediated by DDM1. Disruption of DDM1 compromises this epigenetic complementarity, leading to transcriptomic imbalances that reduce heterosis.

**Highlight:** Loss of DDM1 reduces biomass heterosis in *Arabidopsis thaliana* hybrids by disrupting DNA methylation and transcriptome stability, revealing a novel salicylic acid–independent epigenetic mechanism.

## Introduction

Heterosis, also known as hybrid vigor, refers to the superior performance of the first filial generation (F_1_) plants compared to their parental lines. This effect is particularly evident in certain parental combinations. Heterosis is important in agriculture, where it is widely exploited to enhance agronomic traits such as yield, stress resistance, and growth rate (Fujimoto *et al*., 2018). The benefits of heterosis have led to the development of F_1_ hybrid varieties in many crop species, resulting in considerable improvements in agricultural productivity (Fujimoto *et al*., 2018). In *Arabidopsis thaliana*, heterosis is observed in shoot size (leaf area and fresh weight), with notable growth advantages emerging a few days after sowing and continuing throughout the growth period (Fujimoto *et al*., 2012; 2018; Groszmann *et al*., 2014; Meyer *et al*., 2004; 2012).

Various mechanisms, including dominance, overdominance, pseudo-overdominance, and epistasis, have been proposed to explain heterosis from a quantitative genetic perspective. Genetic studies such as quantitative trait locus (QTL) mapping and genome-wide association studies (GWAS) have been used to identify the genetic factors underlying heterosis (Fujimoto *et al*., 2018; Wu *et al*., 2021). QTLs associated with traits like whole leaf area and shoot dry biomass, which are believed to contribute to heterosis, have been identified in *A. thaliana* (Meyer *et al*., 2010). While a correlation between the genetic distance of parental lines and the degree of heterosis has been observed, this correlation is not particularly strong in *A. thaliana* (Meyer *et al*., 2004; Vasseur *et al*., 2019; Yang *et al*., 2017,), suggesting that heterozygosity in specific genomic regions, rather than overall genetic distance of parental lines, plays a key role in the manifestation of heterosis (Fujimoto *et al*., 2018; Wu *et al*., 2021).

In addition to genetic factors, recent studies have highlighted the importance of epigenetic regulation in heterosis (Miyaji and Fujimoto, 2020, Nishimura *et al*., 2025; Shiraki *et al*., 2023,). Epigenetics refers to mechanisms that influence gene regulatory networks without altering the underlying DNA sequence (Fujimoto *et al*., 2012). Notably, although this phenomenon is restricted to certain epigenetic recombinant inbred lines (epiRILs) that exhibit variation in DNA methylation and share identical or almost identical genetic backgrounds, an epiHybrid, a cross between an epiRIL and the wild type, has shown heterosis despite both parental lines being genetically identical (Dapp *et al*., 2015; Lauss *et al*., 2018). This suggests that epigenetic differences between parental lines can contribute to heterosis, supporting the hypothesis that epigenetic regulation, in addition to genetic factors, contributes to the enhanced performance observed in hybrid progeny. Furthermore, the loss-of-function mutation of DECREASE IN DNA METHYLATION 1 (DDM1), a chromatin remodeling factor, has been shown to diminish heterosis in *A. thaliana* (Kawanabe *et al*., 2016; Zhang *et al*., 2016a), further emphasizing the role of chromatin regulators and epigenetic mechanisms in the expression of heterosis (Kakoulidou and Johannes, 2024).

The multifunctional plant hormone, salicylic acid (SA), plays a critical role as bio-stimulator and regulator of plant development (Rivas-San *et al*., 2011; Groszmann *et al*., 2015). SA is mediated by two distinct pathways: the isochorismate (IC) pathway and the phenylalanine ammonia-lyase (PAL) pathway, both occurring in plastids. Among these, the IC pathway is the primary route for pathogen-induced SA synthesis in *A. thaliana* (Klessig *et al*., 2018; Kumar, 2014; Rekhter *et al*., 2019). SA is synthesized from chorismate by ISOCHORISMATE SYNTHASE 1 (ICS), and SA produced via this pathway is essential for both local (LAR) and systemic (SAR) acquired resistance responses (Wildermuth *et al*., 2001). Many autoimmune mutants show SA over-accumulation and exhibit dwarfism (Janda and Ruelland, 2015; Van Wersch *et al*., 2016). Mutation of SA biosynthetic enzymes such as in the *salicylic acid induction deficient 2* (*sid2*) mutant, or disruption of SA accumulation (e.g., by introducing the *NahG* transgene), results in increased biomass and seed yield (Abreu *et al*., 2009). It has been suggested that SA concentration levels have a significant effect on heterosis, possibly by influencing the growth-defense balance (Groszmann *et al*., 2015; Miller *et al*., 2015; Yang *et al*., 2017; Zhang *et al*., 2016a).

In this study, we used a *ddm1* mutant to investigate the relationship between epigenetics and early seedling biomass heterosis in hybrids between the C24 and Columbia-0 (Col) accessions of *A. thaliana*. By combining RNA-sequencing (RNA-seq) and whole genome bisulfite sequencing (WGBS), we conducted a comprehensive analysis of gene expression and DNA methylation levels to explore the potential molecular mechanisms underlying heterosis. Additionally, by using mutants defective in SA biosynthesis, we examined the relationship between SA and heterosis.

## Materials and methods

### Plant materials and growth condition

Columbia-0 (Col) and C24 accessions were used as parental lines. Mutant lines of *ddm1-1* (Col) (Vongs *et al*., 1993), *ddm1-9* (C24) (Chang *et al*., 2005), *eds1-2* (Col) (Bartsch *et al*., 2006), and *pad4-1* (Col) (Jirage *et al*., 1999) were used. The NahG transgenic line expressing bacterial salicylate hydroxylase in a Col background was used. We crossed *eds1-2* (Col) or *pad4-1* (Col) with C24, and *eds1-2* or *pad4-1* mutants introgressed into C24 were obtained from BC_4_F_2_ populations. Plants were grown in a controlled environment under a 16-h/8-h light/dark cycle at 22 °C. Seeds were sown on plastic dishes containing Murashige and Skoog agar medium supplemented with 1.0% sucrose (pH 5.7), and seedlings were transferred to soil at 14 days after sowing (DAS). Primer sequences used for genotyping the *ddm1-1*, *ddm1-9*, *eds1-2*, *pad4-1* and NahG lines are shown in Table S1.

### Measuring rosette diameter, leaf area, and salicylic acid contents

Rosette diameter and leaf area were measured for evaluation of plant size or heterosis. Rosette diameter is defined as the maximum diameter of the rosette, measured between the two largest leaves at a specific developmental stage. Rosette diameter depends on the length of the leaf blade and petiole. The leaf area of the entire plant was calculated from photographs using image analysis with Image-J software (http://rsb.info.nih.gov/ij/)

SA contents were determined by liquid chromatography-tandem mass spectrometry (LC-MS/MS), as previously described (Mori *et al*., 2025). Briefly, approximately 100 mg of aerial tissue collected at 14 days after sowing (DAS) was flash-frozen and homogenized in an extraction solvent consisting of 80% acetonitrile and 1% acetic acid, supplemented with isotope-labeled internal standards including D4-SA (OlChemim, Olomouc, Czech Republic). The detailed procedures for sample purification and mass spectrometry conditions were based on previously published methods (Gupta *et al*., 2017), with minor modifications. The extracts were sequentially purified using Oasis HLB, MCX, and WAX columns (Waters Corporation, Milford, MA, USA). For SA analysis, after eluting the main acidic fraction, SA was specifically eluted from the Oasis WAX column using 3% formic acid in 97% acetonitrile. Quantification was performed on a 6410 Triple Quad LC/MS system (Agilent Technologies, Santa Clara, CA, USA) equipped with a CAPCELLPAC ADME HR S2 analytical column (OSAKASODA, Osaka, Japan) and an XDB-C8 guard column (Agilent Technologies).

### RNA extraction, RNA-sequencing, and gene ontology analysis

Total RNA was isolated from aerial tissues at 14 DAS in Col, C24, and C24 x Col hybrids of both wild type and *ddm1* mutant backgrounds using the SV Total RNA Isolation System (Promega). The TruSeq RNA Sample Preparation Kit was used to prepare sequence libraries for RNA-sequencing, following the manufacturer’s protocol (TruSeq RNA Library Prep Kit v2, Illumina, San Diego, CA, USA). Sequencing was performed using the Illumina HiSeq2500 System with 100bp paired-end reads. Low-quality reads were filtered using FaQCs version 2.10 (Lo and Chain, 2014), and the filtered reads were aligned to the *A. thaliana* reference genome (TAIR10) (Lamesch *et al*., 2012) using Tophat2 (Kim *et al*., 2013). Gene expression levels were quantified as fragments per kilobase per million (FPKM) using Cuffdiff v2.2.1 (Trapnell *et al*., 2012).

Gene ontology (GO) enrichment analysis was performed using the agriGO tool (Du *et al*., 2010). The background reference for RNA-seq was the list of genes that displayed expression above background levels in either the parental or F_1_ samples (Fujimoto *et al*., 2012). Statistical tests for the enrichment of functional terms used the hypergeometric test and false discovery rate (FDR) correction for multiple testing at a threshold of 5% FDR.

### DNA extraction and whole genome bisulfite sequencing

Genomic DNA from aerial tissues at 14 DAS in Col, C24, and C24 x Col hybrids of both wild type and *ddm1* mutants was isolated by using DNeasy Plant Mini Kit (Qiagen). 100 ng of DNA was bisulfite converted using the EZ DNA Methylation-Gold kit (Zymo Research), and a sequence library was constructed according to the post-bisulfite adaptor tagging (PBAT) protocol (Miura and Ito, 2015). PBAT libraries were sequenced using Illumina HiSeq 2500 sequencing system (paired-end, 100bp).

WGBS reads were mapped to the *A. thaliana* reference genome (TAIR10) using Bowtie2 version 2.4.4 and Bismark v0.23.1. Methylation levels of CG, CHG, and CHH contexts were calculated based on the numbers of methylated and unmethylated reads at each cytosine position, using the bismark_methylation_extractor script with the paired-end option. The methylation level at each cytosine position was calculated as the ratio of methylated cytosine reads to the total number of reads covering that site. Differentially methylated regions (DMRs) were identified using the R package methylKit (v1.18.0). Differentially methylated genes (DMGs) were defined as genes containing DMRs within gene bodies (exon and intron) or 200 bp upstream or downstream of the gene.

### DNA methylation analysis using methylKit

WGBS data were processed using the R package methylKit (v1.18.0) for the quantification and comparative analysis of DNA methylation levels. The input files consisted of Bismark-generated coverage files for each sample, representing cytosine methylation status across the genome. A total of 12 samples, representing wild-type (WT) and *ddm1* mutant backgrounds from multiple genotypes (Col, C24, and F_1_) and two biological replicates, were analyzed.

Coverage files were imported using the methRead function with the pipeline=“bismarkCoverage” option. Cytosine sites with extremely low coverage (read count < 5) and excessively high coverage (top 0.1% quantile) were filtered out using filterByCoverage function. Read coverage normalization across samples was performed using the normalizeCoverage function with the “median” method.

After unifying methylation calls across all samples using unite function, methylation values were pooled by condition (e.g., “Col-WT”, “Col-ddm1”, etc.) using the pool function. The genome was segmented into 500 bp sliding windows with a step size of 250 bp using tileMethylCounts function, retaining only those windows containing at least 20 covered cytosines.

Tiled methylation data were reorganized into treatment groups using the reorganize function to enable pairwise comparisons. DMRs were identified using the calculateDiffMeth function with the “fast.fisher” test and Benjamini-Hochberg (BH) correction for multiple testing.

## Results

### Decrease heterosis was caused by the loss of DDM1 function

Since DNA methylation changes are inherited in subsequent generations, crosses between *DDM1*/*ddm1* heterozygotes of C24 and Col will be the first in which the loss of DDM1 function manifests in the F_1_ generation. The F_1_ plants derived from a cross between a heterozygote for the *ddm1-9* mutation in C24 (*DDM1*/*ddm1-9*) and a heterozygote for the *ddm1-1* mutation in Col (*DDM1*/*ddm1-1*) were produced, and rosette diameter of the F_1_s was measured. From 14 to 28 DAS, F_1_ plants homozygous for *ddm1* showed a smaller rosette diameter than those heterozygous for *ddm1* or homozygous for *DDM1* plants (Fig.1A).

**Fig. 1.**
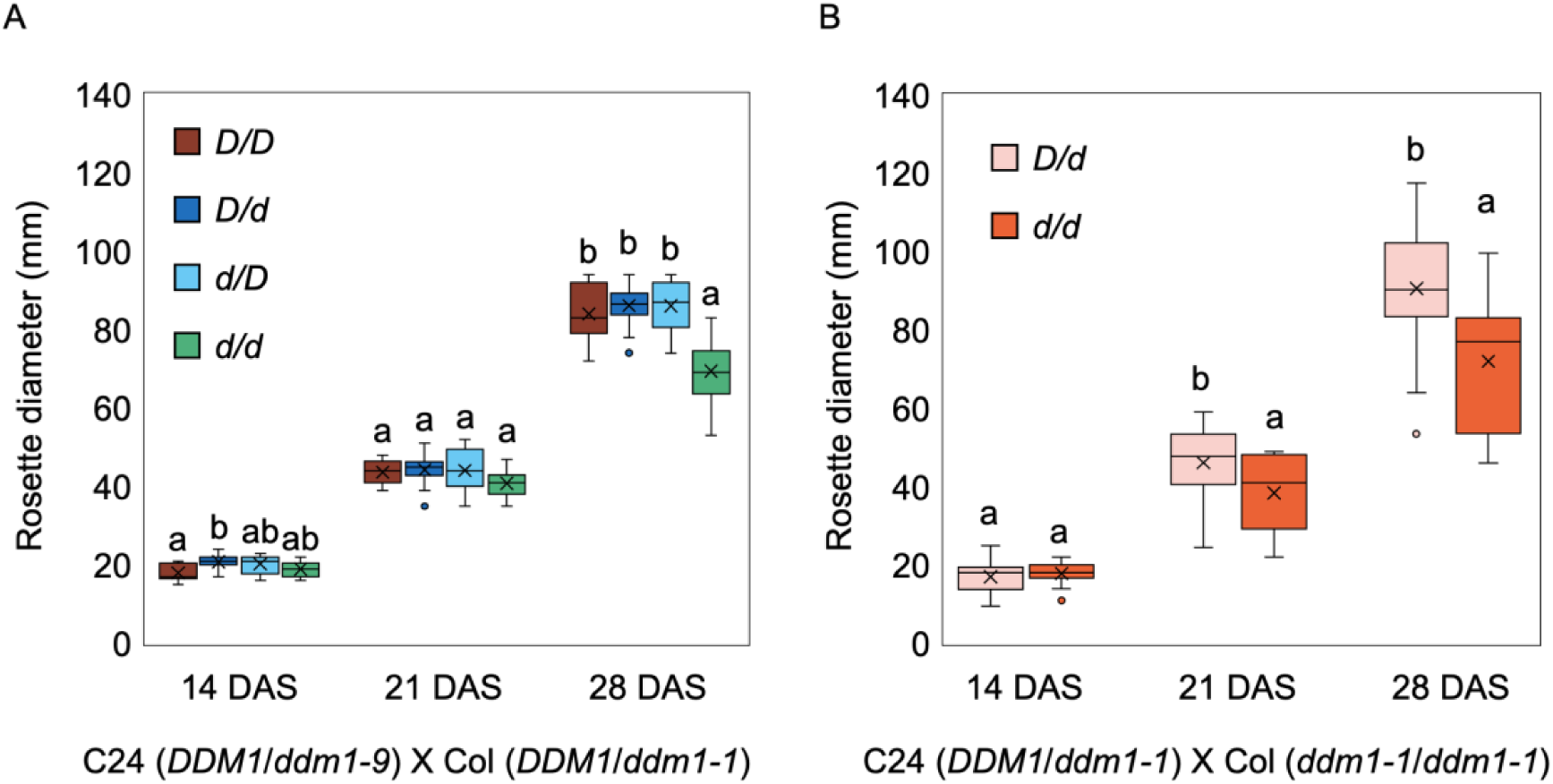
Rosette diameter in hybrids between heterozygous of *DDM1*/*ddm1* in C24 and in Col (A), and hybrids between *ddm1* backcrossed plants in C24 and *ddm1-1* in Col (B). Different letters indicate significant differences [*p* < 0.05 Tukey’s test (A), Student’s *t*-test (B)]

We also generated a BC_4_F_1_ population with a heterozygous *ddm1-1* mutation; *ddm1-1* in the Col background was crossed with C24 and subsequently backcrossed with C24 while maintaining the *ddm1-1* mutation. The F_1_ was obtained by crossing BC_4_F_1_ plants carrying a heterozygous mutation (*DDM1*/*ddm1-1*) with the *ddm1* mutant in Col (*ddm1-1/ ddm1-1*), F_1_ plants homozygous for *ddm1* exhibited a smaller rosette diameter to those heterozygous for *ddm1* (Fig. 1B).

### Genes differentially expressed between wild type and the ddm1 mutant in the F_1_ and its parental lines

RNA-seq analysis was performed using aerial tissues from wild-type plants (Col, C24, C24xCol) and *ddm1* mutants (*ddm1*-Col, *ddm1*-C24, *ddm1*-C24x*ddm1*-Col) at 14 DAS. Approximately 13.0 million to 16.3 million reads (about 91.3 to 98.5% of total reads) were uniquely mapped to the reference genome (Table S2). Differentially expressed genes (DEGs) were identified using an FDR of 5%. Between wild type and the *ddm1* mutant, 1,948 DEGs (1,615 up-regulated and 333 down-regulated in *ddm1*) in Col, 1,357 DEGs (1,132 up-regulated and 225 down-regulated in *ddm1*) in C24, and 1,836 DEGs (1,494 up-regulated and 342 down-regulated in *ddm1*) in F_1_ were identified (Table 1). 402 were upregulated and 34 genes were downregulated in both F_1_ and parental lines (Fig. 2). 427 genes were uniquely upregulated, and 201 genes were uniquely downregulated in the *ddm1*-F_1_, referred to as F_1_-specific upregulated genes (F_1_ up-d) and F_1_-specific downregulated genes (F_1_ down-d), respectively, in response to the *ddm1* mutation (Fig. 2). Approximately 50% of upregulated genes in *ddm1* were classified as transposable elements (TEs) in Col, C24, and F_1_, whereas less than 2% of downregulated genes in *ddm1* were classified as TEs (Table 1).

**Fig. 2.**
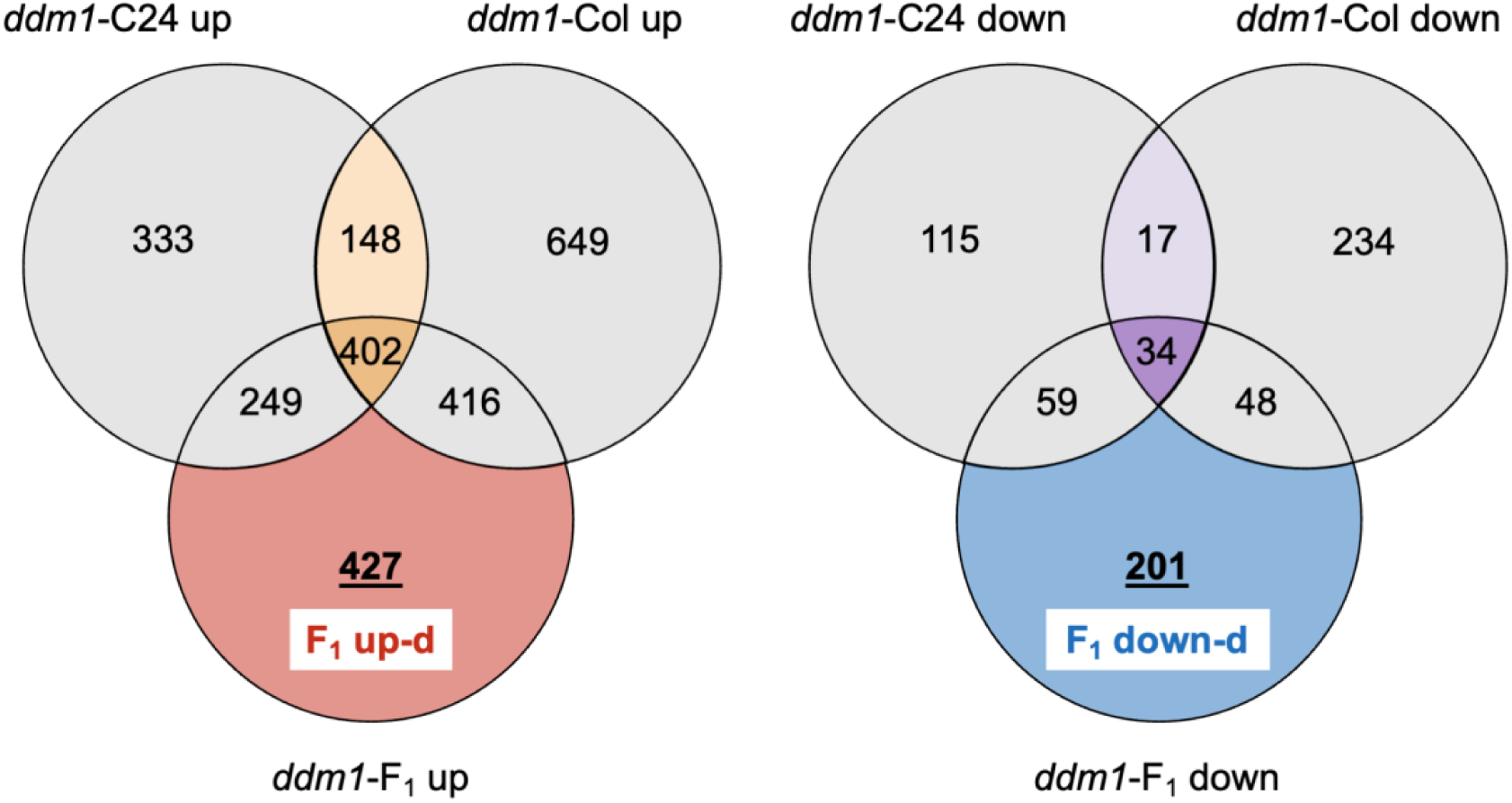
Venn diagram of differentially expressed genes between *ddm1* mutant and wild type.

**Table 1.** Number of differentially expressed genes between wild type and *ddm1* mutant.

|  |  |  |  |  |  |  |  |  |  |
| --- | --- | --- | --- | --- | --- | --- | --- | --- | --- |
|  | *ddm1*-Col vs. Col |  |  | *ddm1*-C24 vs. C24 |  |  | *ddm1*-F<sub>1</sub> vs. F<sub>1</sub> |  |  |
|  | *ddm1*>WT | *ddm1*<WT | Total | *ddm1*>WT | *ddm1*<WT | Total | *ddm1*>WT | *ddm1*<WT | Total |
| Number of differentially expressed genes <sup>a</sup> | 1,615 | 333 | 1,948 | 1,132 | 225 | 1,357 | 1,494 | 342 | 1,836 |
| Transposable elements <sup>b</sup> | 805 | 5 | 890 | 565 | 2 | 567 | 852 | 3 | 855 |
| Percentage (b/a) | 49.85% | 1.50% | 45.69% | 49.91% | 0.89% | 41.78% | 57.03% | 0.88% | 46.57% |

|  | *ddm1*-Col vs. Col |  |  |  |  |
| --- | --- | --- | --- | --- | --- |
|  | *ddm1*>WT | *ddm1*<WT | Total |  |  |
| Number of differentially expressed genes <sup>a</sup> | 1,615 | 333 | 1,948 |  |  |
| Transposable elements <sup>b</sup> | 805 | 5 | 890 |  |  |
| Percentage (b/a) | 49.85% | 1.50% | 45.69% |  |  |
| *ddm1*-C24 vs. C24 |  |  | *ddm1*-F<sub>1</sub> vs. F<sub>1</sub> |  |  |
| *ddm1*>WT | *ddm1*<WT | Total | *ddm1*>WT | *ddm1*<WT | Total |
| 1,132 | 225 | 1,357 | 1,494 | 342 | 1,836 |
| 565 | 2 | 567 | 852 | 3 | 855 |
| 49.91% | 0.89% | 41.78% | 57.03% | 0.88% | 46.57% |

The up and downregulated genes in the *ddm1* mutants of Col, C24, and F_1_ were categorized into GO cellular component (CC), GO molecular function (MF), and GO biological process (BP). A total of 26, 65, and 8 categories were overrepresented in upregulated genes of *ddm1*-Col, *ddm1*-C24, and *ddm1*-F_1_, respectively, with only one GO category shared between Col and F_1_ (Fig. S1, Table S3, S4, S5). Specifically, ‘plastid part’ and ‘chloroplast part’ GO terms were overrepresented in the upregulated genes of *ddm1*-Col, while ‘response to abiotic stimulus’ and ‘response to biotic stimulus’ GO terms were overrepresented in the upregulated genes of *ddm1*-C24 (Fig. 3). In the upregulated genes of *ddm1*-F_1_, the GO terms, ‘cell wall’ and ‘xyloglucan:xyloglucosyl transferase activity’, were overrepresented (Fig. 3). Of the 33 *XYLOGLUCAN ENDOTRANSGLUCOSYLASE/HYDROLASE* (*XTH*s) genes, one, five, and nine were expressed at higher levels in wild-type plants compared to *ddm1* mutants of Col, C24, and F_1_, respectively; of the three lines, *XTH*s expression levels tended to be higher in *ddm1*-F_1_(Fig. S2).

**Fig. 3.**
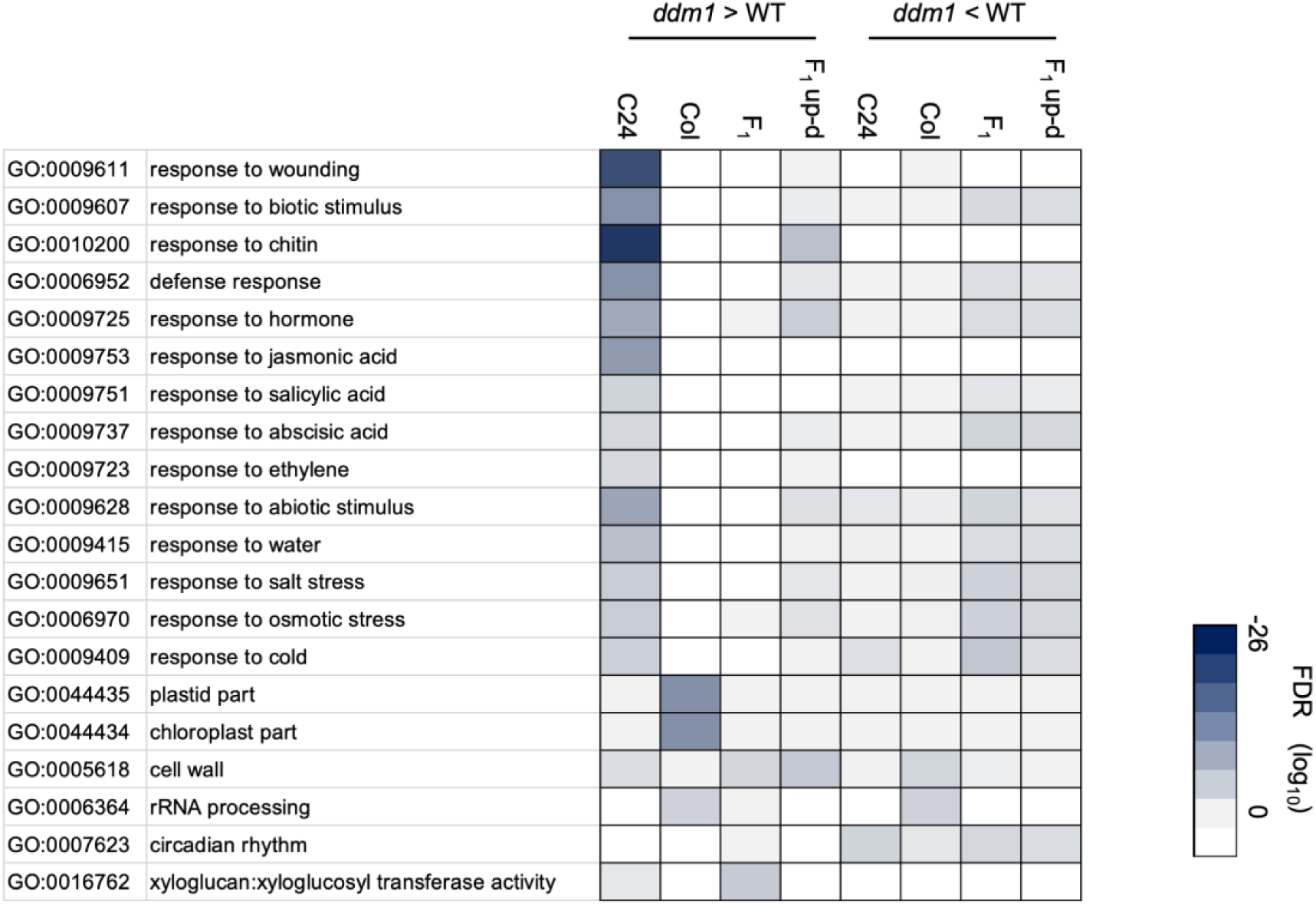
GO terms overrepresented in up and downregulated genes in the *ddm1* mutants of Col, C24, and F_1_. Log_10_ values of FDR are shown with the scale from light gray to dark blue. White column represents not listed by GO analysis by agriGO.

In the *ddm1* mutants of Col, C24, and F_1_, 9, 3, and 22 categories, respectively, were overrepresented in the downregulated genes, with only two GO categories shared between C24 and F_1_ (Fig. S1, Tables S6, S7, S8). Specifically, the GO terms, ‘cell wall’ and ‘rRNA processing’, were overrepresented in the downregulated genes of *ddm1*-Col, while ‘circadian rhythm’ was overrepresented in downregulated genes of *ddm1*-C24 (Fig. 3). In the downregulated genes of *ddm1*-F_1_, the GO terms, ‘response to abiotic stimulus’, ‘response to biotic stimulus’, and ‘circadian rhythm’, were overrepresented (Fig. 3).

Thirty-five and seventeen categories were overrepresented in the F_1_ up-d and F_1_ down-d genes, respectively (Tables S9, S10). Specifically, genes categorized as ‘response to chitin’, ‘cell wall’, and ‘response to hormone’ were overrepresented in F_1_ up-d genes, while genes categorized as ‘response to salt stress’, ‘response to osmotic stress’, ‘response to abscisic acid’ and ‘response to water’ were overrepresented in F_1_ down-d genes (Fig. 3).

### Identification of differentially expressed genes between parental lines and their F_1_

In wild-type plants, 2,536 DEGs were identified between F_1_ and Col (F_1_>Col, 1,229; F_1_<Col, 1,307), and 1,765 DEGs between F_1_ and C24 (F_1_>C24, 945; F_1_<C24, 820) (Table S11). In *ddm1* mutants, 1,357 DEGs were identified between *ddm1*-F_1_ and *ddm1*-Col (*ddm1*-F_1_>*ddm1*-Col, 696; *ddm1*-F_1_<*ddm1*-Col, 661), and 2,270 DEGs between *ddm1*-F_1_ and *ddm1*-C24 (*ddm1*-F_1_>*ddm1*-C24, 1,358; *ddm1*-F_1_<*ddm1*-C24, 912) (Table S11). 227 and 194 DEGs overlapped between F_1_>Col and *ddm1*-F_1_>*ddm1*-Col and between F_1_<Col and *ddm1*-F_1_<*ddm1*-Col, respectively (Fig. S3). 429 and 438 DEGs overlapped between F_1_>C24 and *ddm1*-F_1_>*ddm1*-C24 and between F_1_<C24 and *ddm1*-F_1_<*ddm1*-C24, respectively (Fig. S3). Genes that showed higher expression levels in *ddm1*-F_1_ than in either *ddm1*-Col or *ddm1*-C24, but not in F_1_ compared to Col or C24, were defined as *ddm1*-F_1_ specific upregulated genes (dF_1_sp-up) (Fig. S3). Using the same criteria, we defined *ddm1*-F_1_ specific downregulated genes (dF_1_sp-down), wild-type F_1_-specific upregulated genes (wtF_1_sp-up), and wild-type F_1_-specific downregulated genes (wtF_1_sp-down) (Fig. S3).

The dF_1_sp-up, dF_1_sp-down, wtF_1_sp-up, wtF_1_sp-down genes were categorized based on GO annotations, and 38, 13, 3, and 0 GO terms were found to be overrepresented, respectively (FDR < 0.001) (Table S12, 13, 14). GO terms such as ‘response to chitin’, ‘xyloglucan metabolic process’, ‘response to abiotic stimulus’, ‘cell wall’, ‘ADP binding’, and ‘response to cold’ were overrepresented in dF_1_sp-up (Fig. 4). GO terms including ‘oxidoreductase activity’, ‘ADP binding’, ‘cell periphery’ and ‘transcription corepressor activity’ were overrepresented in dF_1_sp-down (Fig. 4). GO terms such as ‘ADP binding’, ‘transcription factor activity, sequence-specific DNA binding’, and ‘cell periphery’ were overrepresented in wtF_1_sp-up (Fig. 4).

**Fig. 4.**
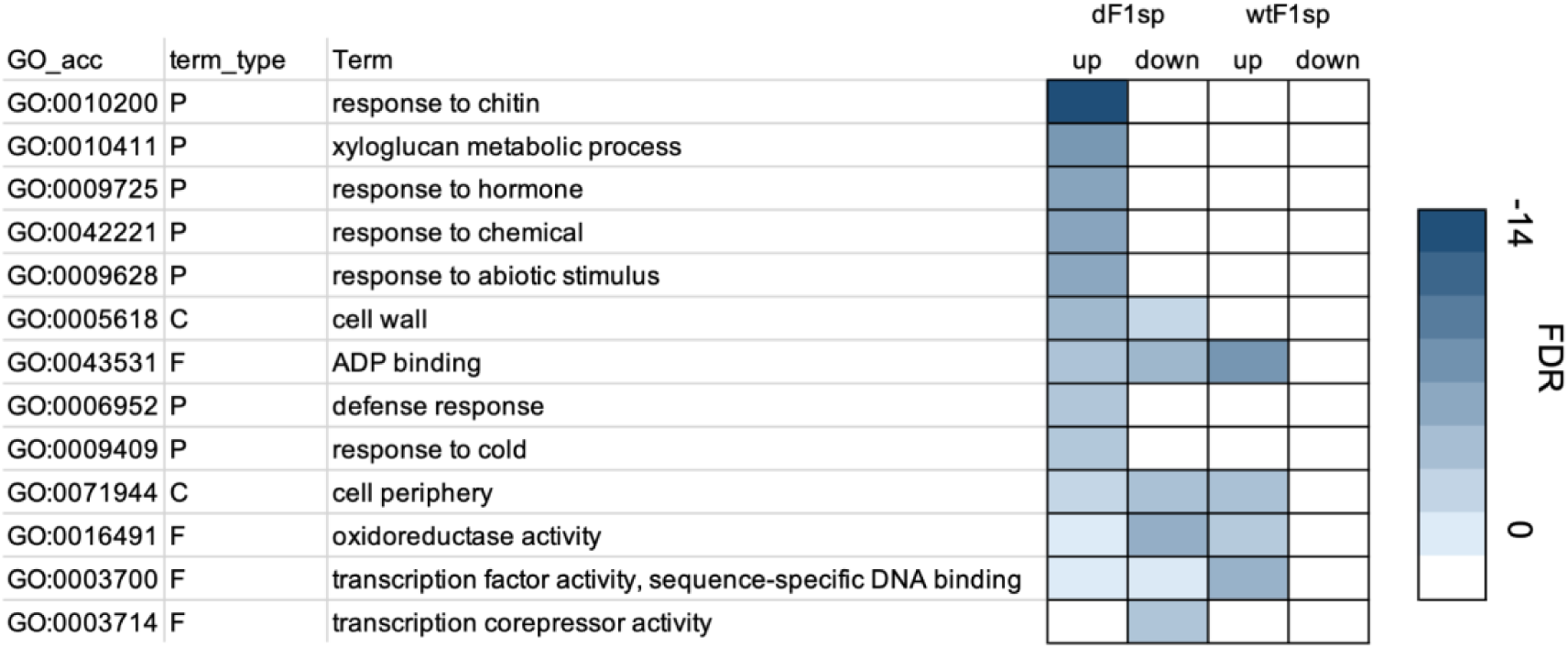
GO terms overrepresented in dF_1_sp-up, dF_1_sp-down, wtF_1_sp-up, and wtF1sp-down genes. Log_10_ values of FDR are shown with the scale from light blue to dark blue. White column represents not listed by GO analysis by agriGO.

### Identification of differentially methylated genes between wild type and ddm1 mutants in F_1_ *and its parental lines*

We performed WGBS on aerial tissues of wild type (Col, C24, C24xCol) and *ddm1* mutants (*ddm1*-Col, *ddm1*-C24, *ddm1*-C24x*ddm1*-Col) at 14 DAS (Table S15). Between Col and *ddm1*-Col, we identified 5,904, 4,827, and 23,898 DMRs in the CG, CHG, and CHH contexts, respectively; 95%, 93%, and 76% of these were hypomethylated in *ddm1*-Col (Table S16). There were 5,780, 4,874, and 22,489 DMRs in the CG, CHG, and CHH contexts, respectively, between C24 and *ddm1*-C24; 99%, 92%, and 67% of these were hypomethylated in *ddm1*-C24 (Table S16). There were 6,131, 5,488, and 25,526 DMRs in the CG, CHG, and CHH contexts, respectively, between F_1_ and *ddm1*-F_1_; 98%, 82%, and 72% of these were hypomethylated in *ddm1*-F_1_ (Table S16). Genes whose genic regions overlapped with DMRs were designated as DMGs. There were 1,251, 352, and 1,391 DMGs in the CG, CHG, and CHH contexts, respectively, between Col and *ddm1*-Col; 81%, 58%, and 73% of DMGs were hypomethylated in *ddm1*-Col (Table S17). There were 1,156, 421, and 1,290 DMGs in the CG, CHG, and CHH contexts, respectively, between C24 and *ddm1*-C24; 95%, 39%, and 58% of DMGs were hypomethylated in *ddm1*-C24 (Table S17). There were 1,343, 746, and 1,921 DMGs in the CG, CHG, and CHH contexts, respectively, between F_1_ and *ddm1*-F_1_; 91%, 29%, and 64% of DMGs were hypomethylated in *ddm1*-F_1_ (Table S17). Approximately 8% of upregulated genes in *ddm1* mutants overlapped with hypo-DMGs (Fig. 5), while other combinations of DEGs and DMGs showed even lower overlap percentages (Fig. S4).

**Fig. 5.**
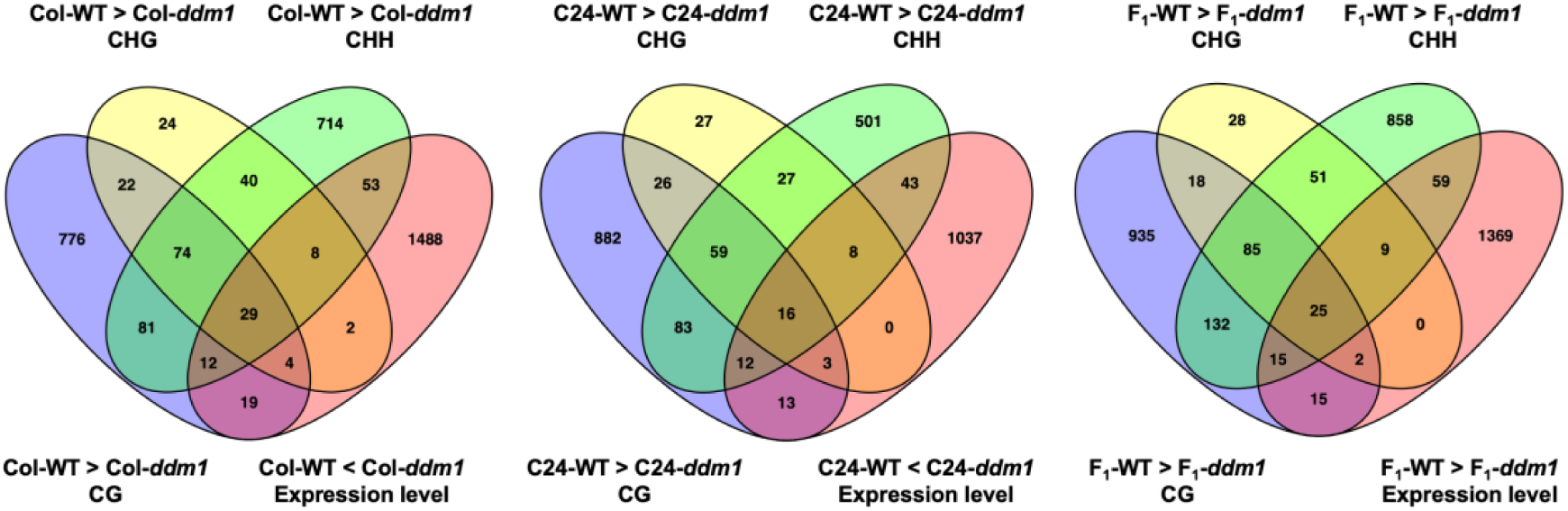
Venn diagram of hypomethylated genes and upregulated genes in *ddm1* mutants.

### Identification of differentially methylated regions between parental lines and their F_1_

The DMRs were identified among the three wild type and *ddm1* mutant lines: 3,222-5,904 in the CG context, 1,094-4,827 in the CHG context, and 7,506-23,898 in the CHH context (Table S18). In both wild type and *ddm1*, CG methylation tended to be lower in the F_1_ than in Col, whereas CHH methylation tended to be higher in F_1_ than in Col. In the wild-type background, CG methylation tended to be higher in F_1_ than in C24, whereas CHH methylation tended to be lower. In the *ddm1* background, both CG and CHH methylation levels in F_1_ tended to be lower than in C24 (Table S18). We identified 468-1,327 DMGs in the CG context, 174-651 DMGs in the CHG context, and 947-1,609 DMGs in the CHH context (Table S19). Only 1.6%-5.9 % of DEGs overlapped with DMGs (Figs. S5-S10)

### Intraspecific differences in endogenous SA contents are not associated with heterosis

Previously, C24 was shown to have a higher SA content than Col and F_1_, and the SA content in *ddm1*-Col, *ddm1*-C24, and *ddm1*-F_1_ was more than in the corresponding wild-type plants (Zhang *et al.,* 2016a). Zhang *et al*. (2016a) suggested that the F_1_ had an optimal endogenous SA concentration, which led to biomass heterosis. Increased SA content resulted in higher expression levels of genes involved in SA biosynthesis, its activation, downstream SA signaling, and SA-responsive genes (Zhang *et al.,* 2016a). As reported previously, the expression of genes involved in SA biosynthesis and downstream of SA signaling was higher in C24 and *ddm1*-C24 than in other lines, but there was no significant difference between C24 and *ddm1*-C24, except for *PR5* (Fig. 6), which contrasts with the previous findings (Zhang *et al.,* 2016a).

**Fig. 6.**
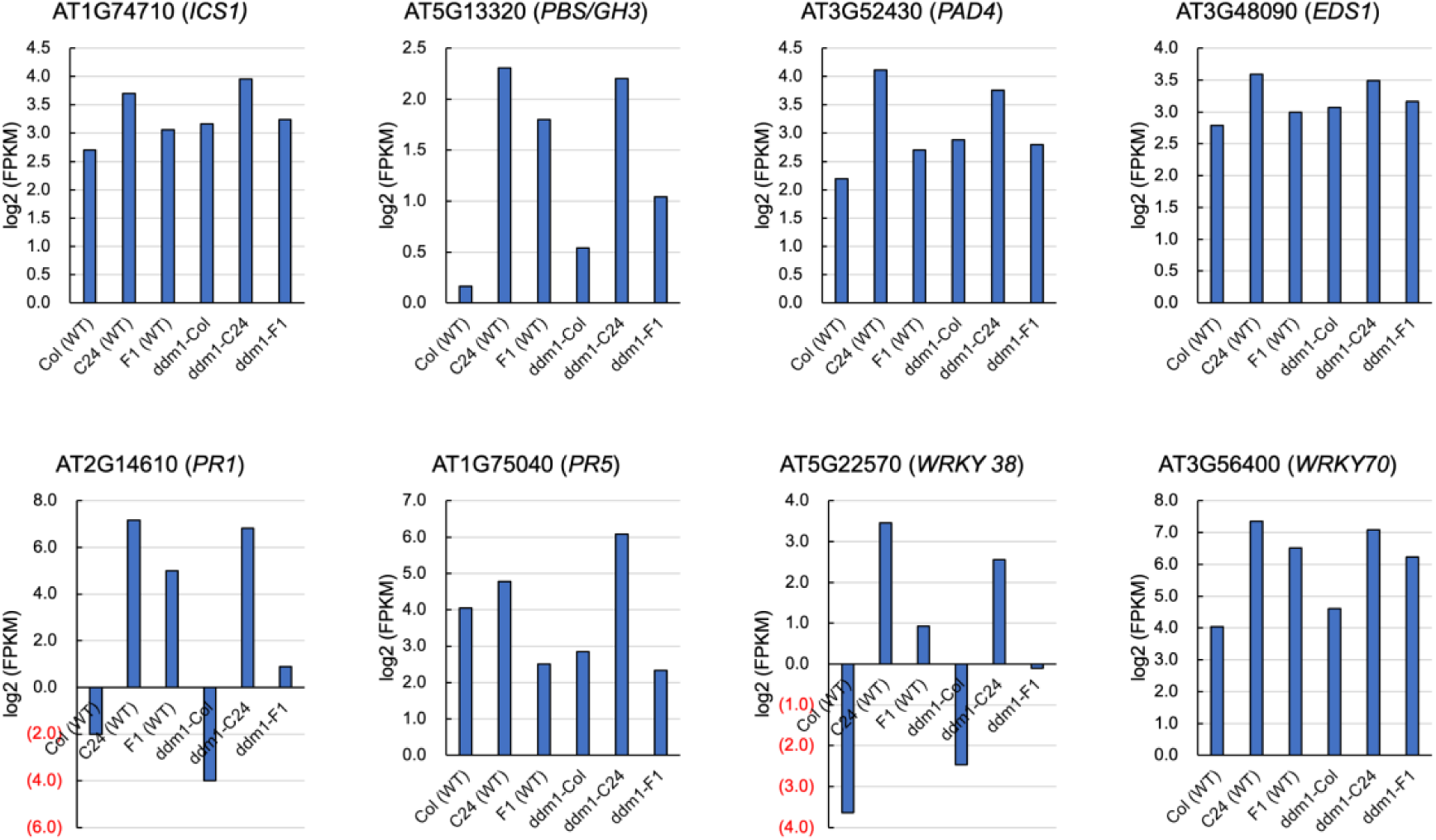
Expression levels of SA-related genes. FPKM, Fragments per kilobase per million

The SA content in C24 was higher than in Col and F_1_ (Fig. 7). To determine whether variation in endogenous SA concentration is important for heterosis, we produced *pad4* and *eds1* mutants in C24 background by backcrossing. In the C24 *pad4* mutant, SA content was lower than in C24 but was still the highest among the three lines (Fig. 7). In the C24 *eds1* mutant, SA content was lower than in C24, and the difference in SA content among the three lines was smaller (Fig. 7). F_1_ hybrids carrying the *pad4* or *eds1* mutation were developed, and both *eds1*-F_1_ and *pad4*-F_1_ showed similar levels of heterosis to F_1_ (Fig. 7, Fig. S11). Although increased plant size in F_1_ between Col_NahG and C24 compared to the F_1_ between Col and C24 was reported (Groszmann *et al*., 2015), our study did not observe a clear increase in growth vigor in F_1_ between C24 and Col_NahG (Fig. S12). These results suggest that endogenous SA content under normal growth conditions is not associated with biomass heterosis.

**Fig. 7.**
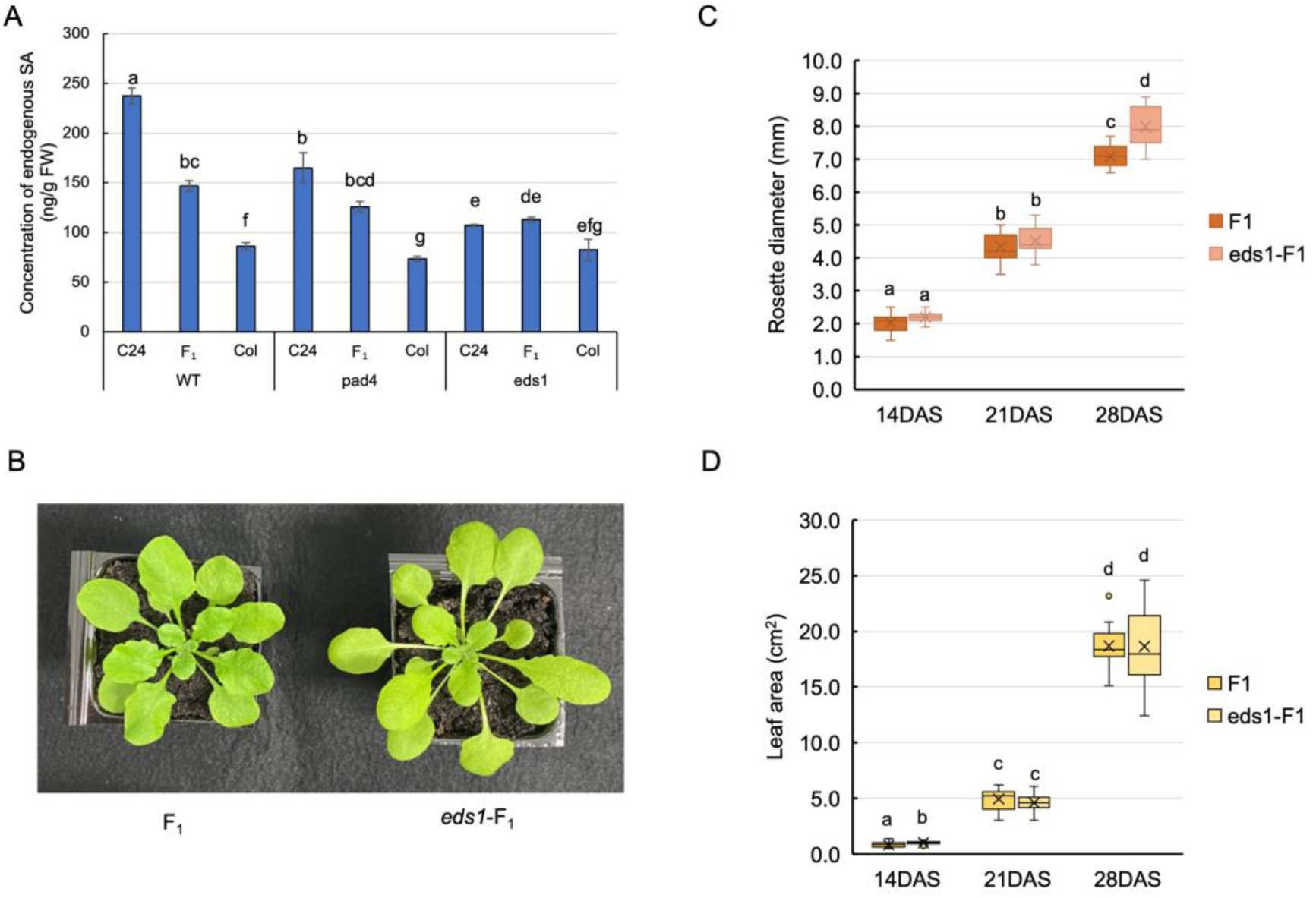
The relationship between SA content and heterosis. (A) Concentration of endogenous SA (B) Plant phenotype of F_1_ and *eds*-F_1_ at 28 days after sowing (DAS) (C) Rosetter diameter of F_1_ and *eds*-F_1_ at 14, 21, and 28 DAS. (D) Leaf area of F_1_ and *eds*-F_1_ at 14, 21, and 28 DAS. Letters indicate the significant differences at *p* < 0.05 (Tukey-Kramer test).

## Discussion

Previous studies have demonstrated that the loss of DDM1 function leads to a reduction in biomass heterosis (Kawanabe *et al*., 2016, Zhang *et al*., 2016a). However, one possibility that could not be ruled out is that the reduction in heterosis was caused by parental plants that had already lost DDM1 function and were hypomethylated. To address this, we specifically designed crosses using *DDM1*/*ddm1* heterozygous parents, ensuring that the loss of DDM1 function would first take place in the F_1_ generation. Despite this design, a decrease in heterosis was still observed. While Zhang et al. (2016a) employed multiple *ddm1* mutant alleles, we used identical *ddm1* alleles in both the Col and C24 backgrounds. Consistently, a reduction in biomass heterosis was observed in our study. These findings strongly suggest that the loss of DDM1 function, even when occurring *de novo* in the F_1_ generation, directly contributes to the attenuation of heterosis. Based on our results and previous studies, we conclude that the reduction in heterosis upon loss of DDM1 function is a robust and reproducible phenomenon.

As described above, DDM1 plays a crucial role in the manifestation of heterosis, however, the underlying molecular mechanisms by which heterosis is attenuated in the absence of DDM1 remain unclear. To address this, we analyzed DNA methylation and gene expression profiles in wild-type and *ddm1*-F_1_ hybrids derived from C24 and Col accessions. A striking finding was the genotype-specific impact of DDM1 loss; while DDM1 dysfunction caused significant transcriptome alterations, the affected genes differed among the Col, C24, and F_1_ genetic backgrounds. This highlights the complex role of DDM1 in regulating gene expression and the potential for epigenetic reprogramming in hybrid plants. Among these differences, a particularly notable observation was the upregulation of *XTH* genes in *ddm1*-F_1_ plants. XTHs are enzymes involved in remodeling plant cell walls by cleaving and rejoining xyloglucan molecules, thereby affecting cell wall loosening and cell expansion (Fry *et al*., 2004; Majda *et al*., 2018). Upregulation of *XTHs* has been observed in heterotic hybrids and hybrid mimic lines (Groszmann *et al*., 2015; Wang *et al*., 2019). In contrast, in our study, expression levels of *XTHs* tended to be higher in *ddm1*-F_1_, specifically *XTH17*, *XTH18*, *XTH19*, and *XTH33*. *XTH17*, *XTH18*, and *XTH19* are phylogenetically closely related (Vissenberg *et al*., 2005). A previous study showed that overexpressing *XTH19* decreased average plant size (Mehraj *et al*., 2020), suggesting that the upregulation of these *XTH*s in *ddm1*-F_1_ may contribute to reduced plant growth and, consequently, reduced heterosis.

Another key insight comes from the downregulation of genes categorized into **‘**circadian rhythm’ in *ddm1*-F₁, even though all samples were harvested at the same time (six hours after lights on, Zeiteber time 6). Previous studies have suggested that a relationship between vegetative growth heterosis and circadian regulation, and CCA1, a central circadian oscillator, plays a role in vegetative growth heterosis (Ni *et al*., 2009; Yang *et al*., 2021). In our dataset, expression level of *CCA1* was higher in *ddm1*-F_1_ compared to wild-type F_1_. It has been shown that increasing or overexpressing the expression level of *CCA1* at noon leads to a reduction in plant size (Ni *et al*., 2009). In addition, this upregulation of *CCA1* in *ddm1*-F_1_ may be related to downregulation of genes categorized into **‘**circadian rhythm’ and could be involved in the attenuation of heterosis in *ddm1*-F_1_ plants. Transcriptome comparison between F_1_ and parental lines revealed differences between the wild-type and *ddm1*-F₁; *ddm1*-F₁ showed upregulation of genes related to abiotic and biotic stress responses in addition to the xyloglucan metabolic process, whereas wild-type F_1_ exhibited increased expression of genes associated with transcriptional regulation. Although many studies have reported altered stress-related gene expression in wild-type F_1_ hybrids compared to parents, the elevated expression of stress-related genes in *ddm1*-F_1_ may be involved in the reduction in the level of heterosis.

Several lines of evidence have suggested a potential involvement of endogenous SA levels in heterosis (Groszmann *et al*., 2015; Zhang *et al*., 2016a). One such observation is that mutants defective in SA biosynthesis often exhibit increased plant size (Janda and Ruelland, 2015; Van Wersch *et al*., 2016), implying a possible trade-off between defense and growth or yield (Yang *et al*., 2021). Furthermore, lower endogenous SA concentrations and lower expression levels of SA biosynthetic genes in F₁ hybrids compared to their mid-parent values (MPV), as well as the enhanced growth observed in NahG-containing C24 plants, have led to the hypothesis that SA metabolism may play a critical role in the manifestation of heterosis (Groszmann *et al*., 2015). In our study, we also found that the SA levels and expression levels of SA-related genes in F₁ hybrids were lower than the MPV. However, this result warrants careful interpretation. The apparent lower levels in F₁ hybrids were primarily driven by the high level in the C24 parent, and the F₁ did not consistently exhibit lower values than Col. In addition, there is also an argument that the optimal concentration of SA is important for heterosis. Under normal conditions, SA levels are low, and the difference between Col and heterotic F₁ hybrid is minimal (Zhang *et al*., 2016a), possibly within the range that could be easily affected by environmental factors as SA content can vary substantially in response to pathogen infection. The F₁ hybrid between Col and Sei accessions exhibits both defense heterosis and growth heterosis (Yang *et al*., 2021). Upon pathogen infection, SA levels increase markedly, with the F₁ showing the highest accumulation. To further examine the relationship between SA metabolism and heterosis, we generated SA-deficient mutants and NahG-containing F₁ plants. These lines exhibited comparable levels of heterosis to wild-type F₁ hybrids. Moreover, although the SA biosynthesis levels were similarly reduced across all three *eds1* mutants, only the F₁ hybrid exhibited heterosis. These findings suggest that SA metabolic capacity alone is not sufficient to explain the occurrence of heterosis, and further investigation is needed to clarify the potential link between SA metabolism and heterosis.

Since DDM1 is involved in maintenance DNA methylation, one possible explanation for the reduction in heterosis is that changes in DNA methylation are responsible. In this study, we examined the relationship between DNA methylation and gene expression, however, the overlap between DMGs and DEGs was limited; only 8% of genes upregulated in *ddm1* mutants compared to wild type showed hypomethylation in *ddm1* mutants. Similar results, indicating a weak correlation between hypomethylation and gene activation, have been reported in *met1* mutants (Srikant *et al*., 2022) and in the *mddcc* mutant, which is completely unmethylated (He *et al*., 2022). These observations suggest that relatively few genes exhibit expression changes directly attributable to alterations in DNA methylation. In this study, we were unable to identify specific genes whose expression changes due to reduced DNA methylation were directly responsible for the attenuation of heterosis. A previous study that performed WGBS in *ddm1* mutants of Col, C24, and their F_1_ hybrids found that SA related non-additively expressed genes, which was the focus of this study, were either low or unmethylated in wild-type plants (Zhang *et al*., 2016a). Thus, the relationship between SA-related nonadditive gene expression and DNA methylation has not been found, and it has been suggested that alterations in histone modifications, rather than DNA methylation, may underlie the reduction in heterosis level observed in *ddm1* mutants (Zhang *et al*., 2016a). The role of DNA methylation in heterosis could be further clarified by examining mutants of DNA methyltransferases that are directly involved in DNA methylation.

F_1_ hybrids generally inherit DNA methylation states from both parents; the majority of DNA methylation states in F_1_ hybrids show an additive pattern, while methylation remodeling events such as trans-chromosomal methylation (TCM) and trans-chromosomal demethylation (TCdM) were observed. These methylation remodeling events are thought to be driven by small RNAs (Greaves *et al*., 2016; Zhang *et al*., 2016b), but heterosis still occurs in mutants defective in small RNA biosynthesis (Kawanabe *et al*., 2016; Zhang *et al*., 2016a), suggesting that the additive inheritance of parental DNA methylation states in F_1_ plays a more central role in the establishment of heterosis than DNA methylation remodeling. There was an association between parental pericentromeric DMRs and leaf area heterosis in epiHybrids, suggesting that parental DMRs in these regions are important for heterosis (Kakoulidou *et al*., 2024). Since DNA methylation in centromeric or pericentromeric regions is significantly decreased in *ddm1* mutants, it is possible that the loss of DDM1 function eliminates differences in DNA methylation states between parental lines in these regions. This loss of epigenetic divergence may lead to the reduced heterosis level observed in *ddm1*-F_1_. This finding aligns with the hypothesis that maintaining parental epigenetic divergence in F_1_ is important for the expression of heterosis.

Taken together, our results suggest that the maintenance of parental epigenetic marks is critical for the manifestation of heterosis. In *ddm1*-F₁ hybrids, the disruption of DDM1 function likely compromises the inheritance and maintenance of these epigenetic differences, particularly DNA methylation patterns in key genomic regions such as pericentromeric domains. This loss of parental epigenetic divergence may lead to aberrant gene expression patterns, including dysregulation of stress responses, circadian rhythms, and cell wall remodeling, ultimately resulting in a reduction of heterosis. Thus, the attenuation of heterosis observed in *ddm1* mutants is most plausibly explained by the breakdown of epigenetic complementarity between parental genomes.

## Supplementary Data

**Figure S1-S12:** Figure S1. Venn diagram of the number of overrepresented Gene Ontology (GO) terms. Figure S2. Expression levels of *xyloglucan endotransglucosylase/hydrolases* (*XTH*s). Figure S3. Venn diagram of the number of differentially expressed genes among Col, C24, and F_1_ in wild type and *ddm1* mutants. Figure S4. Venn diagram of the number of differentially methylated genes and differentially expressed genes between wild type and *ddm1* mutants in Col, C24, and F_1._ Figure S5. Venn diagram of the number of differentially methylated genes and differentially expressed gene between Col and C24. Figure S6. Venn diagram of the number of differentially methylated genes and differentially expressed gene between Col and F_1_. Figure S7. Venn diagram of the number of differentially methylated genes and differentially expressed gene between C24 and F_1_. Figure S8. Venn diagram of the number of differentially methylated genes and differentially expressed gene between *ddm1*-Col and *ddm1*-C24. Figure S9. Venn diagram of the number of differentially methylated genes and differentially expressed gene between *ddm1*-Col and *ddm1*-F_1_. Figure S10. Venn diagram of the number of differentially methylated genes and differentially expressed gene between *ddm1*-C24 and *ddm1*-F_1_. Figure S11. Time course pf rosette diameter in pad4 mutant hybrids. Figure S12. Time course pf rosette diameter in F_1_ hybrids containing NahG.

**Table S1-17:** Table S1. Sequences of primers used for genotyping. Table S2. Mapped RNA-sequencing reads to reference genome (TAIR10). Table S3. GO terms of genes showing a higher expression level in *ddm1*-Col than in Col. Table S4. GO terms of genes showing a higher expression level in *ddm1*-C24 than in C24. Table S5. GO terms of genes showing a higher expression level in *ddm1*-F1 than in F_1_. Table S6. GO terms of genes showing a lower expression level in *ddm1*-Col than in Col. Table S7. GO terms of genes showing a lower expression level in *ddm1*-C24 than in C24. Table S8. GO terms of genes showing a lower expression level in *ddm1*-F_1_ than in F_1_. Table S9. GO terms of F_1_ up-d genes. Table S10. GO terms of F_1_ down-d genes. Table S11. Number of differentially expressed genes among Col, C24, and its F_1_ in wild type and *ddm1* mutants. Table S12. GO terms of dF_1_sp-up. Table S13. GO terms of dF1sp-down. Table S14. GO terms of wtF1sp-up. Table S15. Mapped whole genome bisulfite sequence reads to reference genome (TAIR10). Table S16. Number of differentially methylated regions between wild type and *ddm1* mutant. Table S17. Number of differentially methylated genes between wild type and *ddm1* mutant. Table S18. Number of differentially methylated regions between Col, C24 and F_1_ in wild type and *ddm1* mutant. Table S19. Number of differentially methylated genes between Col, C24, and F_1_ in wild type and *ddm1* mutants.

## Acknowledgements

We thank Dr. Naomi Miyaji, Ms. Satoko Takada, Ms. Namiko Nishida, Mr. Kohei Kunita, and Ms. Yumiko Arai for their technical assistance.

## Author Contribution

KM, RW, HY, TM, MAA, YK, and KN performed experiments, and ST, KT and MS performed data analysis. AK and HK made the sequence library and performed sequencing. TM, TM, and YI performed LC-MS/MS. MS, ESD, and RF participated in the design of the study. RW, HY, ESD, and RF drafted the manuscript. All the authors have read and approved the final version of the manuscript.

## Data availability statement

DDBJ; accession no. DRA015525

